# A Novel Cisplatin-Based Prodrug Inhibits Lysine Deacetylases, Suppresses Nucleotide Excision Repair, and Overcomes Resistance

**DOI:** 10.1101/2024.10.30.621050

**Authors:** Ya’ara Negev-Korem, Esther Stern, Hadar Golan-Berman, Elisheva Heilbrun, Subhendu Karmakar, Yoram Soroka, Marina Frušić-Zlotkin, Ofer Chen, Hiba Hassanain, Ori Wald, Dan Gibson, Ron Kohen, Sheera Adar

## Abstract

Cisplatin [cis-diamminedichloroplatinum(II)] is a widely used chemotherapeutic agent that induces cytotoxicity primarily through DNA damage, but drug resistance severely limits its efficacy and use. Cisplatin resistance is complex and multifactorial, involving DNA repair via nucleotide excision repair (NER), and overexpression of lysine deacetylases (KDACs), which reduce chromatin accessibility and alter transcription regulation. The combination of cisplatin and KDAC inhibitors has shown promise in improving treatment efficacy. This improved efficacy has been attributed to increased drug sensitivity due to higher chromatin accessibility, however, this hypothesis has not been validated. In this study, we synthesized a novel Pt(IV) derivative, *cct*-[Pt(NH_3_)_2_Cl_2_(VPA)(PhB)] (cPVP), which combines cisplatin and two KDAC inhibitors, phenyl butyrate and valproic acid. This triple-action prodrug enabled the simultaneous targeting of multiple cancer-related pathways. Compared to cisplatin, cPVP exhibited significantly enhanced damage formation and cytotoxicity. High-resolution mapping of cisplatin damage and repair, however, does not attribute the enhanced damage sensitivity to chromatin accessibility, but rather to increased drug uptake and inhibition of nucleotide excision repair. Moreover, cPVP treatment increased survival in a mouse mesothelioma model, and prevented the development of resistance to both cisplatin and itself in cancer cells. Our findings shed new light on the effect of KDAC inhibition on cisplatin treatment, and suggest that cPVP could serve as a promising alternative to cisplatin in the clinic.

## Introduction

Since its approval in 1979, the metallo-drug cisplatin [cis-diamminedichloroplatinum(II)] is a widely used chemotherapeutic agent (1, 2). It is used for multiple tumor types, notably testicular and ovarian cancers (3, 4), but also lung cancer, head and neck cancer, bladder cancer, cervical cancer and others (5, 6). The continued use of cisplatin is greatly restricted by severe dose-limiting side effects due to lack of tumor selectivity (7), and intrinsic or acquired drug resistance limiting cisplatin efficacy (8).

Cisplatin cytotoxicity is mainly attributed to its covalent interaction with DNA strands to form DNA adducts (9, 10). The main mechanism for eliminating cisplatin DNA adducts in human cells is nucleotide excision repair (NER) (11, 12). NER is divided into two sub-pathways that differ in the damage recognition step: global genome repair (GGR), in which repair factors recognize damage directly throughout the genome, and transcription-coupled repair (TCR), in which an RNA polymerase that is blocked by the damage acts as the damage recognition factor, recruiting repair factors to actively transcribed regions. After damage recognition, the two pathways share the subsequent steps of repair, which involve the excision of a single strand oligonucleotide containing the lesion (∼26nt) from the genome, followed by gap filling by DNA polymerase and ligation by a DNA ligase (13–16).

Increased NER plays a role in cisplatin resistance (17, 18), which is complex and multifactorial as it involves numerous cellular pathways and activities (8, 19). Accelerated rate of adduct repair may attenuate the apoptotic process and thus facilitate the survival of resistant cells (11, 17). Another mechanism involved in cisplatin resistance is overexpression of lysine deacetylases (KDACs), which are key epigenetic modifiers (19, 20). By removing acetyl groups from lysine residues within histone tails, KDACs (also termed histone deacetylases, HDACs) are responsible for chromatin condensation, thus limiting access to DNA strands and promoting transcriptional repression (21, 22). Additionally, these enzymes are known to deacetylate lysine residues of non-histone proteins, thus regulating numerous biological processes and widely affecting cellular functioning (23). Aberrant activity of KDACs is known to promote resistance to DNA-targeting agents, including cisplatin, by a mechanism that is not entirely clear (24). An additional mechanism involved in resistance to cisplatin is the constitutive activation of the nuclear factor erythroid 2-related factor 2 / Kelch-like erythroid cell-derived protein with CNC homology associated protein 1 (Nrf2/Keap1) pathway (25, 26), which is responsible for the antioxidant and detoxifying defense system of the cell (27). Resistant cells exploit Nrf2 activation to enhance their survival and eliminate cisplatin (28, 29). Considering the contribution of these three mechanisms to cisplatin resistance, simultaneously inhibiting both KDAC activity and Nrf2/Keap1 pathway could be beneficial for retrieving chemo-sensitivity. HDAC inhibition is also known to impede NER activity (30, 31), thereby reinforcing the potential for overcoming cisplatin resistance by simultaneously targeting multiple cellular pathways.

Pt(IV) derivatives of cisplatin have been developed precisely for the purpose of combining cisplatin treatment with additional anti-cancer drugs. These multi-targeting prodrugs can release cisplatin and two additional bioactive moieties inside the cancer cells, thereby enhancing the chances of overcoming resistance and increasing the therapeutic index. The conversion of four coordinate Pt(II) drugs (e.g. cisplatin) to six coordinate Pt(IV) derivatives involves their oxidation followed by axial modification with bioactive ligands, to form a double/triple-action prodrugs (32, 33). As the original cisplatin, these molecules are likely to also harm normal dividing cells more efficiently. However, these compounds are considered relatively inert and stable outside tumor cells, hence are expected to have an increased bioavailability and reduced side effects compared to the original Pt(II) drug (33, 34). When entering the cell, Pt(IV) prodrugs are reduced and release the Pt(II) parent drug and the bioactive molecules, enabling interactions with their intracellular targets **(Figure 1)** (35). Accordingly, Pt(IV) prodrugs have the potential to evade cisplatin resistance and present improved pharmacology, as the bioactive moieties have the potential to act synergistically and efficiently to eradicate cancer cells (33, 34).

**Figure 1.**
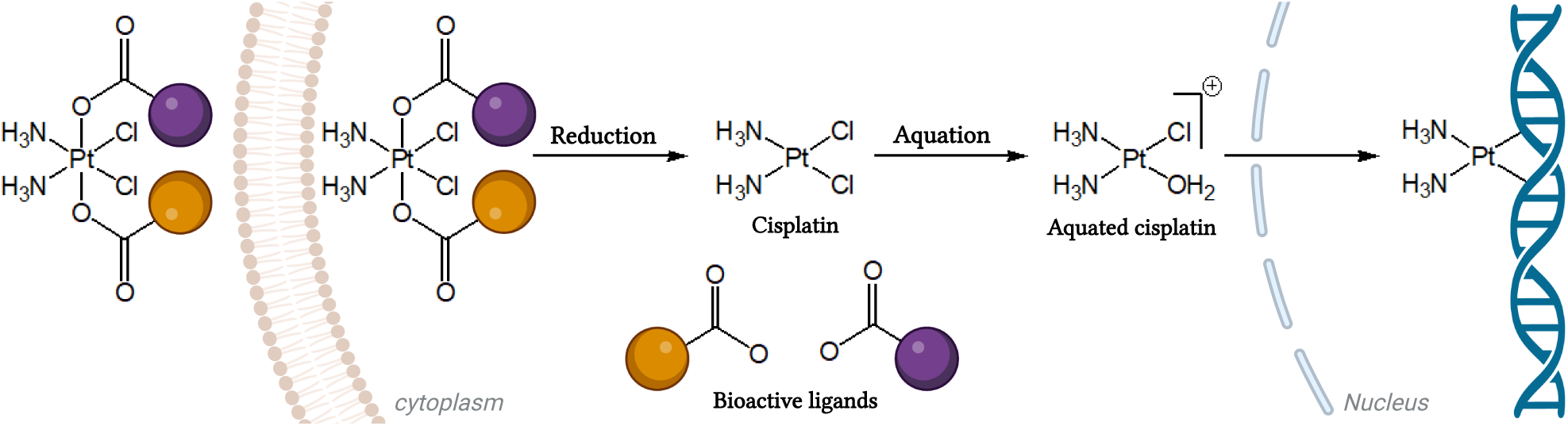
Pt(IV) prodrug. A triple-action Pt(IV) prodrug of cisplatin axially modified by two different bioactive carboxylate ligands (orange and purple). It is inert outside the cell and is activated intracellularly by reduction to the origin moieties. This is followed by aquation of cisplatin to attain DNA covalent binding.

In this study, we have selected 4-phenylbutyric acid (PhB) and valproic acid (VPA) as the bioactive ligands to be conjugated with cisplatin. Both PhB and VPA are KDAC inhibitors (KDAC*i*), thus enabling us to explore the contribution of chromatin accessibility to cisplatin treatment (36, 37). VPA is also an Nrf2 inhibitor (Nrf2*i*), which decreases Nrf2 nuclear content and increases Keap1 expression, providing an additional advantage against cisplatin resistance (38–40). These specific inhibitors were chosen because they possess carboxylic groups, which are essential for conjugation with Pt(IV) (32). Upon reduction, the bond between Pt(IV) and the carboxylate oxygen is broken, releasing the inhibitors in their active form (35). Previously reported dual-action Pt(IV) derivatives of cisplatin with either two PhB or two VPA axial ligands have demonstrated anti-cancer activity and significantly improved cytotoxicity compared to cisplatin (41). However, the combination of both PhB and VPA in a single agent was not reported. Here, we synthesized a novel triple-action Pt(IV) prodrug consisting of PhB and VPA fused to cisplatin, and tested its effect on cisplatin damage formation, repair, cytotoxicity and resistance. Our results indicate KDAC inhibition increases damage-sensitivity in cells, and cPVP holds the potential to overcome cisplatin resistance.

## Results

### Synthesis of a novel triple action prodrug composed from cisplatin, PhB and VPA

The prodrug synthesis started from oxoplatin (*cct*-[Pt(NH_3_)_2_Cl_2_(OH)_2_] the Pt(IV) derivative of cisplatin with two OH axial ligands). The two hydroxido groups present in the axial positions were functionalized via esterification by the anhydride of the corresponding bioactive ligands, in a stepwise manner (**Figure 2**). In brief, oxoplatin was reacted over-night with ∼1.5 equivalents of valproic anhydride in DMSO to yield the intermediate product monovalproato-monohydroxido-oxoplatin, *cct*-[Pt(NH_3_)_2_Cl_2_(VPA)(OH)], which was reacted further with 5 equivalents of 4-phenylbutyric anhydride to yield the desired biscarboxylato product, *cct*-[Pt(NH_3_)_2_Cl_2_(VPA)(PhB)] (cPVP). The progress of the reactions was monitored by ^195^Pt NMR, and the HPLC purified final product was characterized by ^1^H, ^13^C & ^195^Pt NMR spectroscopy, as well as by ESI-MS and elemental analysis (**Supplemental Figure S1A-E**). cPVP was stable after storage at −20°C for at least 3 years (**supplemental Figure S1F**).

**Figure 2.**
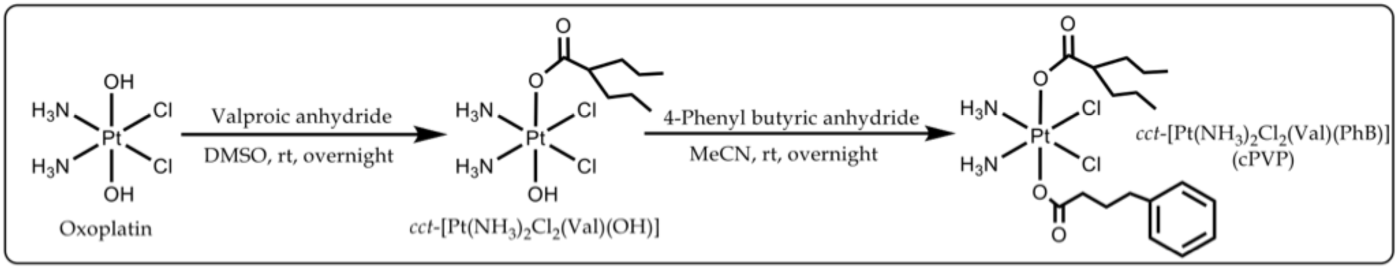
Schematic representation of cPVP synthesis. Cisplatin was oxidized by H_2_O_2_ to oxoplatin, which was successfully carboxylated on axial positions with the anhydrides of VPA and PhB, in a stepwise manner. rt – room temperature.

### cPVP is a more efficient genotoxic agent than cisplatin

In order to evaluate the efficacy of cPVP as an anticancer agent, we compared its cytotoxic effect to that of cisplatin. A2780 ovarian cancer and A549 lung cancer cell lines were treated with ascending concentrations of cPVP or cisplatin for 72 hours, followed by MTT cell-viability assay and IC_50_ assessment. Previously published IC_50_ values for cisplatin are 0.5 - 1.1 µM for A2780 cells (42, 43) and 3.8 – 5.95 µM for A549 cells (43, 44). For both cell-lines, cPVP was much more toxic than cisplatin (**Figure 3A,B**), with up to three orders of magnitude lower IC_50_ values (**Table 1**). As a control, we also compared the cytotoxic effect of cPVP to co-administration of its components. cPVP was found to be more toxic, as co-administration of its components presented identical cytotoxic profile to that of cisplatin alone (**Supplemental Figure S2**).

**Figure 3.**
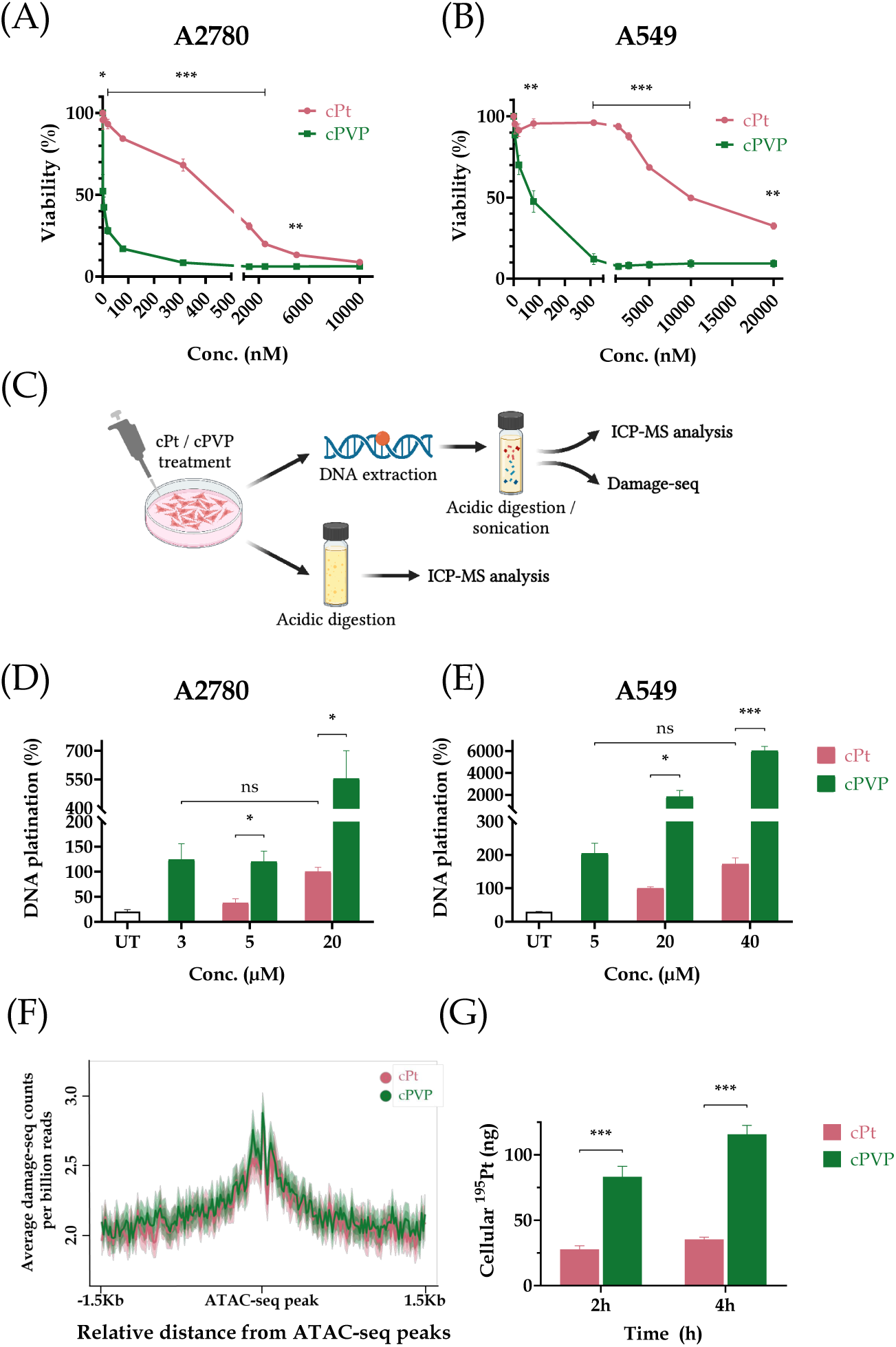
cPVP is a more efficient genotoxic agent than cisplatin. **(A)** Cytotoxicity of cPVP and cisplatin (cPt) in A2780 cells. Cells were incubated for 72 hours with ascending concentrations of the drugs, followed by MTT assay. Data presented as percent of viability normalized to untreated control. Results are based on three biological replicates performed in quadruplets. (**B**) Same as A except data presented for A549 cells. **(C)** Outline of samples preparation for DNA damage assessment, Damage-seq, and cellular uptake evaluation. **(D)** DNA damage formation following cPt and cPVP treatments in A2780 cells treated with cPt or cPVP. Cells were incubated with treatments for two hours before DNA was extracted and subsequently subjected to ICP-MS analysis of ^195^Pt content. The damage caused by 20 µM of cPt was defined as 100% DNA-platination. Results are based on two biological replicates performed in duplicates. (**E**) Same as D except data presented for A549 cells. **(F)** Average Damage-seq signal following cPt and cPVP treatments in accessible regions in A2780 cells. Damage signal is plotted at the 3 Kb regions flanking ATAC-seq peaks from cells treated with either cPt or cPVP. Shadow represents 95% confidence interval for the mean. **(G)** Cellular uptake of cPt (20 µM) and cPVP (3 µM) after two and four hours of incubation in A2780 cells. Treated cells were digested by nitric acid for subsequent ^195^Pt content determination by ICP-MS. Absolute given doses of ^195^Pt: 3,900 ng from 20µM of cPt, and 580 ng from 3µM of cPVP. Data presented as absolute ^195^Pt amounts detected within cells. Results are based on four biological replicates performed in duplicates. All bar graphs indicate mean ± SEM. *P<0.05; **P<0.01; ***P≤0.001 for Student’s *t*-test.

**Table 1:** IC_50_ values of cisplatin and cPVP for A2780 and A549 cell-lines. . Calculated from MTT viability assay after 72 h of incubation with ascending concentrations of the drugs (0 – 10,000 nM for A2780, and 0 – 20,000 nM for A549). Based on three biological replicates performed in quadruplets. Mean ± SEM.

|  | IC<sub>50</sub> |  |
| --- | --- | --- |
|  | cisplatin (nM) | cPVP (nM) |
| A2780 | 582.6 ± 80.6 | 2.21 ± 1.6 |
| A549 | 10124.4 ± 1183.7 | 55.9 ± 17.02 |

Since cisplatin cytotoxicity is mainly attributed to its DNA damaging property (9, 10), we assessed the ability of cPVP to do the same. A2780 and A549 cells were exposed to varying doses of cPVP or cisplatin for two hours followed by DNA extraction. Platinum content in DNA samples was measured by Inductively Coupled Plasma Mass Spectrometry (ICP-MS; **Figure 3C**). Consistent with the higher cytotoxicity, cPVP caused significantly more DNA damage than cisplatin at each concentration (**Figure 3D,E)**.

cPVP requires much lower doses than those of cisplatin both in terms of cytotoxicity and of damage induction. For our subsequent experiments, we sought to treat cells with similar effective doses. Based on DNA damage levels, for the A2780 cell-line, sub lethal doses that elicit damage would be a short (two-hour) incubation with 3 µM cPVP and 20 µM cisplatin, and for A549 cells, 5 µM cPVP vs. 40 µM cisplatin (**Figure 3D,E**). To facilitate accurate comparisons between treatments our follow-up experiments were performed using these ratios as guidelines to obtain similar effective doses.

KDAC inhibition elevates chromatin acetylation and can subsequently increase chromatin accessibility. Thus, KDAC inhibitors could sensitize the genome to damage. Based on genome-wide mapping of cisplatin adducts, however, adducts formation is not strongly affected by chromatin structure (45). To test if the higher damage levels after cPVP treatment reflect enhanced damage formation at accessible chromatin, we mapped cisplatin adducts by Damage-seq in A2780 cells treated with either cisplatin or cPVP for two hours. Accessible regions were determined by ATAC-seq performed under the same conditions (methods and (46)). The higher cisplatin adduct frequency in these accessible regions reflects the elevated frequency of GG di-nucleotides, the main target for cisplatin adducts (**Supplemental Figure S3**). Treatment with cPVP did not alter the damage profiles, which were almost identical for both treatments (**Figure 3F**).

The higher DNA damage induced by cPVP could be due to efficient cellular uptake of the prodrug. To test this, A2780 cells were incubated with drug concentrations that induce similar damage levels (3 µM for cPVP and 20 µM for cisplatin) for two and four hours, and the cellular platinum content was measured using ICP-MS. At each time point, the cellular accumulation of cPVP was greater than that of cisplatin (**Figure 3G**). ^195^Pt content after cPVP treatment revealed significantly higher platinum content associated with the cells compared to cisplatin treatment, despite the lower concentration of cPVP. Moreover, cisplatin exhibited minimal cellular accumulation even after four hours (approximately 1%), while cPVP cellular uptake reached 20% of the initial dose. Thus, the increased cell-associated platinum concentration indicates there is increased cellular uptake of cPVP and supports its enhanced ability to cause DNA damage.

### The PhB and VPA components of cPVP inhibit KDAC and NRF2 activity

To study the effect of cPVP on KDAC activity, we assessed the acetylation levels of histone H3. A2780 cells were cultured for 24 hours with cisplatin, cPVP or bisPhB (*cct*-[Pt(NH_3_)_2_(PhB)_2_Cl_2_]), which is a previously reported Pt(IV) prodrug known to inhibit KDAC activity (41), and served as positive control. Substantially higher acetylation levels of histone H3 were found in the presence of cPVP or bisPhB vs. cisplatin treatment **(Figure 4A,B; Supplemental Figures S4, S5)**. Since KDAC activity regulates chromatin condensation, we employed ATAC-seq to evaluate the effect of cPVP on chromatin accessibility across the genome (47). cPVP treatment resulted in elevated accessibility compared to cisplatin treatment (**Figure 4C**), consistent with KDAC inhibition.

**Figure 4.**
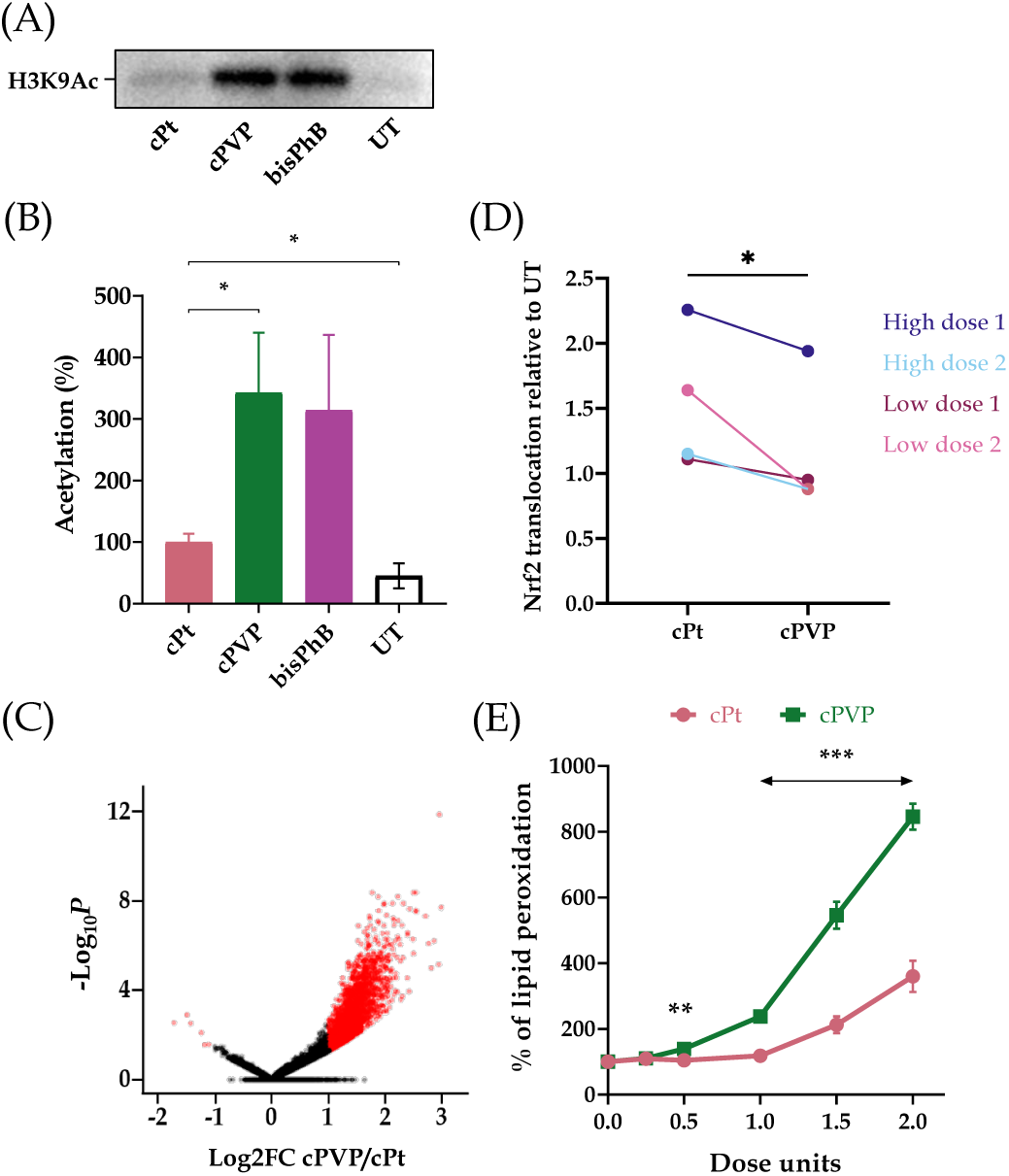
The PhB and VPA components of cPVP inhibit KDAC and NRF2 activity. **(A)** Elevated levels of histone H3 acetylation in the presence of cPVP. A2780 cells were treated with 5 µM of cPVP, 5 µM of bisPhB or 30 µM of cPt for 24 hours. Shown is a representative western blot for H3K9Ac. **(B)** Quantification of western blots from three biological replicates. Acetylation under cPt treatment was defined as 100%. Bands intensity was normalized to total protein. Mean ± SEM. *P<0.05, for one-tailed Student’s *t*-test **(C)** Volcano plot for differential chromatin accessibility at ATAC peaks in cPVP vs cPt treated samples. A2780 cells were treated with either 30 µM of cPVP or 200 µM of cPt for two hours prior to ATAC-seq. **(D)** cPVP reduces Nrf2 translocation to the nucleus relative to cisplatin. A2780 cells were incubated for four hours with either 3 µM of cPVP vs. 20 µM of cPt, or 10 µM of cPVP vs. 70 µM of cPt. Nucleic and cytosolic fractions were extracted and Nrf2 levels were quantified in each. Data expressed as the ratio between treated to untreated nuclei-to-cytoplasm Nrf2 levels ratios. Results are based on two biological replicates for each set of treatments. *P<0.05 for paired Student’s *t*-test. **(E)** cPVP decreases cells’ ability to defend against oxidative stress. A2780 cells were treated with a range of 0 – 2 doses of cPt or cPVP for six hours, followed by over-night ROS induction and TBAR assay. Lipid peroxidation levels were normalized to cells viability using MTT assay. Data presented as percent of untreated control. One dose unit was defined as 20 µM for cPt and 3 µM for cPVP. Results are based on four biological replicates performed in quadruplets. Mean ± SEM. **P<0.01, ***P≤0.001 for Student’s *t*-test.

In order to evaluate the inhibitory effect of cPVP on the Nrf2/Keap1 pathway, we examined its influence on Nrf2 translocation into the nucleus, a crucial step in the pathway’s activation. A2780 cells were incubated with sets of either low or high doses of cPVP and cisplatin for four hours. Next, cytosolic and nuclear fractions were extracted from cells (**Supplemental Figure S6**) and were used for Nrf2 sub-cellular partition quantification. At similar effective doses, cPVP reduces the translocation of Nrf2 into the nucleus relative to cisplatin (**Figure 4D**).

Since Nrf2 activation enhances the antioxidant capacity of the cell, we investigated whether the diminished translocation of Nrf2 to the nucleus following cPVP treatment is reflected in the ability of the cells to defend against oxidative stress. A2780 cells were treated with cisplatin or cPVP for six hours followed by the addition of an oxidative stressor for overnight incubation. The levels of lipid peroxidation products were then measured and normalized to cell viability using the TBAR and MTT assays, respectively. At similar effective doses, cPVP treatment led to a significant increase in lipid peroxidation levels compared to cisplatin treatment (**Figure 4E**), suggesting that cPVP impedes the cells’ protective response against oxidative stress. These results provide further support for Nrf2 inhibition by cPVP.

### cPVP inhibits nucleotide excision repair in cancer cells

To measure repair, we follow the decrease from the initial damage levels with time. Correct measurement of the initial damage levels necessitated the cessation of DNA damage accumulation following the removal of treatments. To validate this, A2780 cells were treated with cisplatin or cPVP for two hours, followed by the removal of treatments. DNA was extracted from cells sampled up to four hours post-treatment, and analyzed for ^195^Pt content using ICP-MS. There was no noteworthy accumulation or reduction of DNA damage at any of the time points for both treatments (**Figure 5A**), indicating the damage levels measured after two hours of treatment and drug removal reflect the initial damage levels in repair experiments.

**Figure 5.**
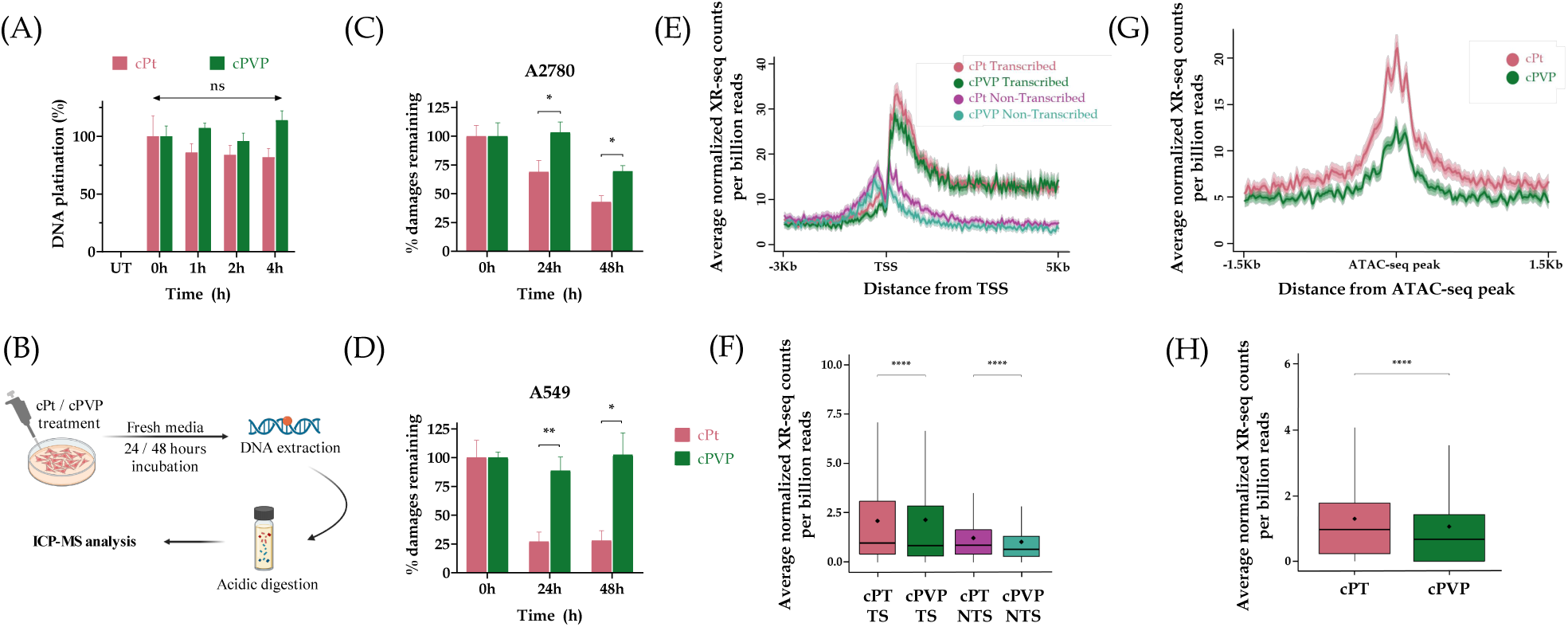
cPVP inhibits nucleotide excision repair in cancer cells. **(A)** DNA damage accumulation post-treatment. A2780 cells were treated with 3 µM of cPVP or 20 µM of cPt for two hours prior to treatments removal. DNA was extracted after 0, 1, 2 and 4 hours post-treatments and subjected to ^195^Pt content determination by ICP-MS. The initial damage measured at time of wash (0h) was defined as 100% DNA-platination for each drug. Results are based on four biological replicates performed in duplicates. Mean ± SEM. ns - not significant for Student’s *t*-test compared to initial damage from each drug. **(B)** Outline of DNA-repair kinetics assay. **(C)** DNA repair kinetics following cPt and cPVP treatments in A2780 cells treated with 3 µM of cPVP or 20 µM of cPt. Drug-containing media was replaced after two hours of incubation and DNA was extracted at 0, 24 and 48 hours post-treatment for ICP-MS analysis. The initial damage was defined as 100% for each drug. Results are based on two biological replicates performed in duplicates. Mean ± SEM. *P<0.05, **P<0.01 for Student’s *t*-test. **(D**) Same as C except A549 cells were treated with 5 µM of cPVP or 40 µM of cPt. **(E)** Average XR-seq repair signal normalized to Damage-seq signal over annotated genes plotted 3 Kb upstream and 5 Kb downstream of the transcription start site (TSS, n=6597). Signal is plotted separately for the transcribed and non-transcribed strands. A2780 cells were treated with 30 µM of cPVP or 200 µM of cPt for two hours prior to XR-seq libraries preparation. The data represent the average of two biological replicates. Shadow represents 95% confidence interval for the mean. **(F)** Total XR-seq read counts over the same genes in E in the transcribed and non-transcribed strands. **** P<0.0001, based on Wilcoxon signed-rank test with Bonferroni correction. Boxes represent range between 75th and 25th percentile, the line represents the median and the diamond the mean. **(G)** Average XR-seq repair signal normalized to Damage-seq signal at the 3 Kb regions flanking ATAC-seq peaks representing accessible chromatin regions. Average cPt and cPVP repair signals are plotted for 7526 and 12048 ATAC-seq peaks measured in cPt and cPVP treated cells, respectively. A2780 cells were treated with 30 µM of cPVP or 200 µM of cPt for two hours prior to ATAC-seq and three hours prior to XR-seq libraries preparation. The data represent the average of two biological replicates. Shadow represents 95% confidence interval for the mean. **(H)** Total XR-seq read counts over the same ATAC-seq peaks from G. **** P<0.0001, based on Wilcoxon signed-rank test with Bonferroni correction. Boxes represent range between 75th and 25th percentile, the line represents the median and the diamond the mean.

To assess the impact of cPVP on DNA repair, A2780 and A549 cells were treated with similar effective doses of cisplatin or cPVP for two hours. Drug-containing media was replaced with fresh media, and cells were sampled at 0, 24, and 48 hours. DNA was extracted from the cells and subjected to measurement of ^195^Pt content by ICP-MS (**Figure 5B**). In both cell lines, adduct repair in cPVP treated cells was suppressed compared to cisplatin treatment (**Figure 5C,D**). In A2780 cells, no repair was observed 24 hours after cPVP treatment, contrasting with 30% repair observed after cisplatin treatment. Furthermore, in A549 cells, no significant repair was observed even 48 hours following cPVP treatment, opposed to 70% repair after cisplatin treatment. These results indicate a strong inhibition of NER by cPVP.

To elucidate the inhibitory effect of cPVP on NER we performed XR-seq, a unique method for high-resolution genome-wide mapping of the excised oligos released during NER (45). The genome-wide profile of repair was very similar for both damaging treatments. Consistent with our previous cisplatin repair maps, higher repair was measured on the transcribed vs. the non-transcribed strands in genes, indicating a preference for transcription-coupled repair of the cisplatin adducts (45, 48). (**Figure 5E,F**). Still, slightly but significantly lower XR-seq signal was observed in cPVP treatment compared to cisplatin alone. Similarly, analysis of repair at accessible chromatin regions (defined by ATAC-seq peaks measured under the same conditions) showed lower cisplatin repair after cPVP treatment. (**Figure 5G,H**). While the overall pattern of repair was similar, the lower enrichment could reflect lower efficiency of repair in cPVP treatment compared to cisplatin.

### cPVP increases cancer-survival and may prevent resistance

To test the potential of cPVP as an anticancer drug, we used a mouse mesothelioma model. AB12 mesothelioma cells were injected into the peritoneum of BALB/c mice to induce tumor formation. On days 6 and 13, cPVP or vehicle solution were injected to the peritoneum of tumor-bearing mice. cPVP treatment increased the median survival of tumor bearing mice in this model from 31 to 39.5 days, highlighting its potential as an effective chemotherapeutic agent (**Figure 6A**).

**Figure 6.**
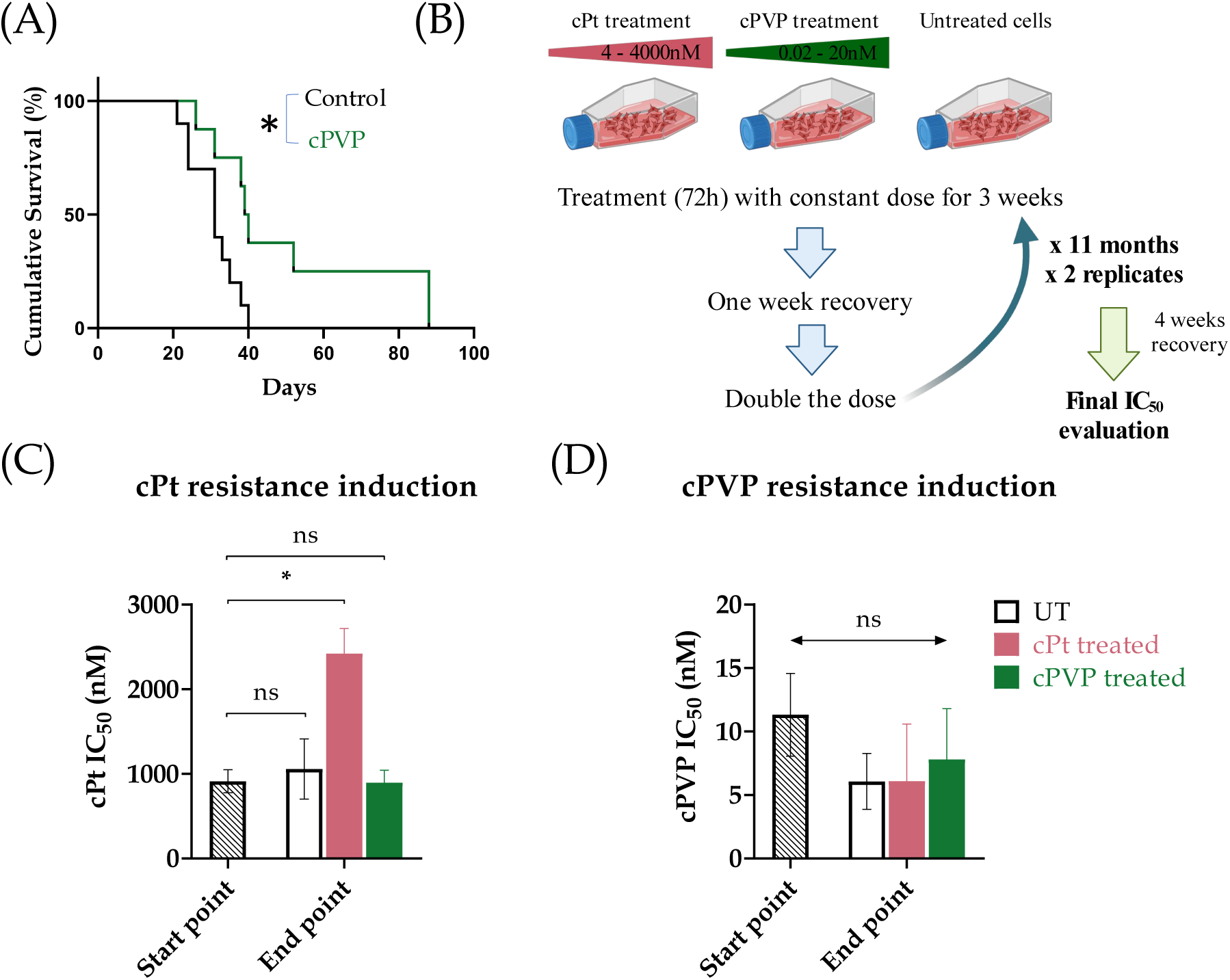
cPVP as a potential anti-cancer drug. **(A)** Cumulative survival of mice injected with AB12 cells to induce tumors, and then injected with either 200mG cPVP or vehicle solution after 6 and 13 days. **(B)** Outline of prolonged resistance induction assay performed on the A2780 cell line. **(C)** IC_50_ values of cisplatin in A2780 cells, before and after long-term exposure to cPt (4 – 4000 nM) or cPVP (0.02 – 20 nM). Results are based on two biological replicates. **(D)** IC_50_ values of cPVP in A2780 cells, before and after long-term exposure to cPt (4 – 4000 nM) or cPVP (0.02 – 20 nM). Results are based on two biological replicates. In all graphs, mean ± SEM are presented. *P<0.05; **P<0.01;

We next assessed the potential of long-term treatment with cPVP to develop resistance. To induce cisplatin or cPVP resistance in A2780 cells, we exposed them to increasing concentrations of the treatments (**Figure 6B**). Initial doses were set to 0.003xIC_50_ of either cPVP or cisplatin. After three weeks of exposure, the doses were doubled. Exposures continued up to doses of 4xIC_50_, requiring almost a year, followed by a four-week period of recovery in which cells were cultured without any treatment. After this long-term treatment with cisplatin, cells exhibited a significant increase in cisplatin IC_50_ value, indicating the development of resistance to the drug. In contrast, cells treated with cPVP presented no change in cisplatin IC_50_ value, indicating that cPVP prevents cells from developing resistance to cisplatin (**Figure 6C**). Moreover, under both conditions there was no change in cPVP IC_50_ values, suggesting that cPVP not only overcomes cisplatin resistance but also prevents the development of self-resistance (**Figure 6D**).

## Discussion

Although broadly used, cisplatin-based chemotherapy is limited due to its severe side effects and multifactorial drug resistance. While the unfavorable toxicity profile of cisplatin (primarily nephrotoxicity and neuropathy) was improved by the development of the second-generation platinum agent, carboplatin (49, 50), overcoming drug resistance was achieved only for colorectal cancers by the development of oxaliplatin, the third-generation platinum drug (51). Yet, the need remains for further platinum drugs that circumvent resistance.

The synthesis of Pt(IV) prodrugs is a well-established approach aimed to improve cisplatin treatment, reduce side effects, and potentially overcome drug resistance (33, 34). Herein, we have utilized this method to synthesize a novel triple-action Pt(IV) prodrug, cPVP, composed from cisplatin, and two KDAC*i*: PhB and VPA, where VPA is also an NRF2*i*. Compared to cisplatin, cPVP demonstrated enhanced efficacy against both cisplatin –sensitive and –resistant cell-lines, as indicated by significantly lower IC_50_ values at the nM scale (as opposed to µM scale for cisplatin), and high DNA-damaging abilities. These findings, together with the extended survival of the mice in the mesothelioma model, suggest that cPVP holds promise as an improved treatment option with lower dosage requirements, and could be potentially effective in treating cancers that are resistant to cisplatin. As for side effects, considering the inertness of Pt(IV) derivatives in terms of binding to proteins and nucleophiles in the blood stream(35), we expect that cPVP will exert less side effects than cisplatin.

The increased cellular uptake of cPVP, attributed to its greater lipophilicity, but only partially accounts for its heightened efficiency. While cPVP kills cells 150-300 times more efficiently than cisplatin, it only induced 3-30-fold more DNA damage. Our results indicate that the PhB and VPA components of the drug functionally contribute to this enhanced efficiency. Since PhB and VPA are negatively charged in physiological fluids, they do not penetrate the negatively charged cell membranes resulting in *in-vitro* IC50 values in the mM range (37, 52). The conjugation of PhB and VPA to Pt(IV) neutralizes the negative charge of these ligands and increases the lipophilicity compared to cisplatin, resulting in efficient cellular accumulation, significantly decreasing the IC_50_ of cPVP to the nM range. Regarding KDAC activity, the high acetylation of histone H3 and elevated chromatin accessibility provide evidence for the inhibitory effect of the cPVP components on KDACs (**Figure 4A-C**). As a prodrug of the Nrf2-inducer cisplatin, and the Nrf2-inhibitor VPA, cPVP possesses the potential for two counter-effects on the Nrf2/Keap1 pathway. Our results indicate that the inhibitory effect of VPA on Nrf2 is stronger than the inductive effect of the cisplatin component. While cisplatin induces Nrf2 translocation to the nucleus, this translocation was reduced after cPVP exposure (**Figure 4D**). Similarly, the ability of the cells to defend against oxidative stress was significantly diminished under cPVP treatment (**Figure 4E**).

We previously reported that chromatin structure affects cisplatin repair but not damage formation, suggesting that KDACs contribution to cisplatin resistance is through influencing NER efficiency (45). Since cPVP caused alterations in chromatin condensation, we assessed its effect on DNA repair by NER. DNA repair was substantially inhibited in the cell lines post exposure to cPVP. These results suggest that one or both of the cPVP components – PhB and/or VPA, impede DNA repair, and consequently cell survival, after treatment. Given that the genomic patterns of repair were largely similar after cPVP and cisplatin treatment, it is not likely that cPVP inhibits repair indirectly through its effects on chromatin structure. The ability of KDACs to deacetylate non-histone proteins, including NER factors (53), may take part in cisplatin resistance (54). For that reason, the KDAC*i* properties of cPVP grant it a wider pharmacological potential beyond “triple-action”.

Though numerous Pt(IV) derivatives have been previously reported, the assessment of their potential to overcome drug resistance was mainly based on IC_50_ values in cisplatin-resistant cell-lines (34). In the case of the novel prodrug cPVP, we conducted a prolonged resistant study that emphasized cPVP’s ability to not only overcome existing cisplatin-resistance, but also to prevent cells from developing resistance to itself or to cisplatin. We attribute these outcomes to the multiple parallel negative effects that cPVP exerts on cancerous cells (**Figure 7**).

**Figure 7.**
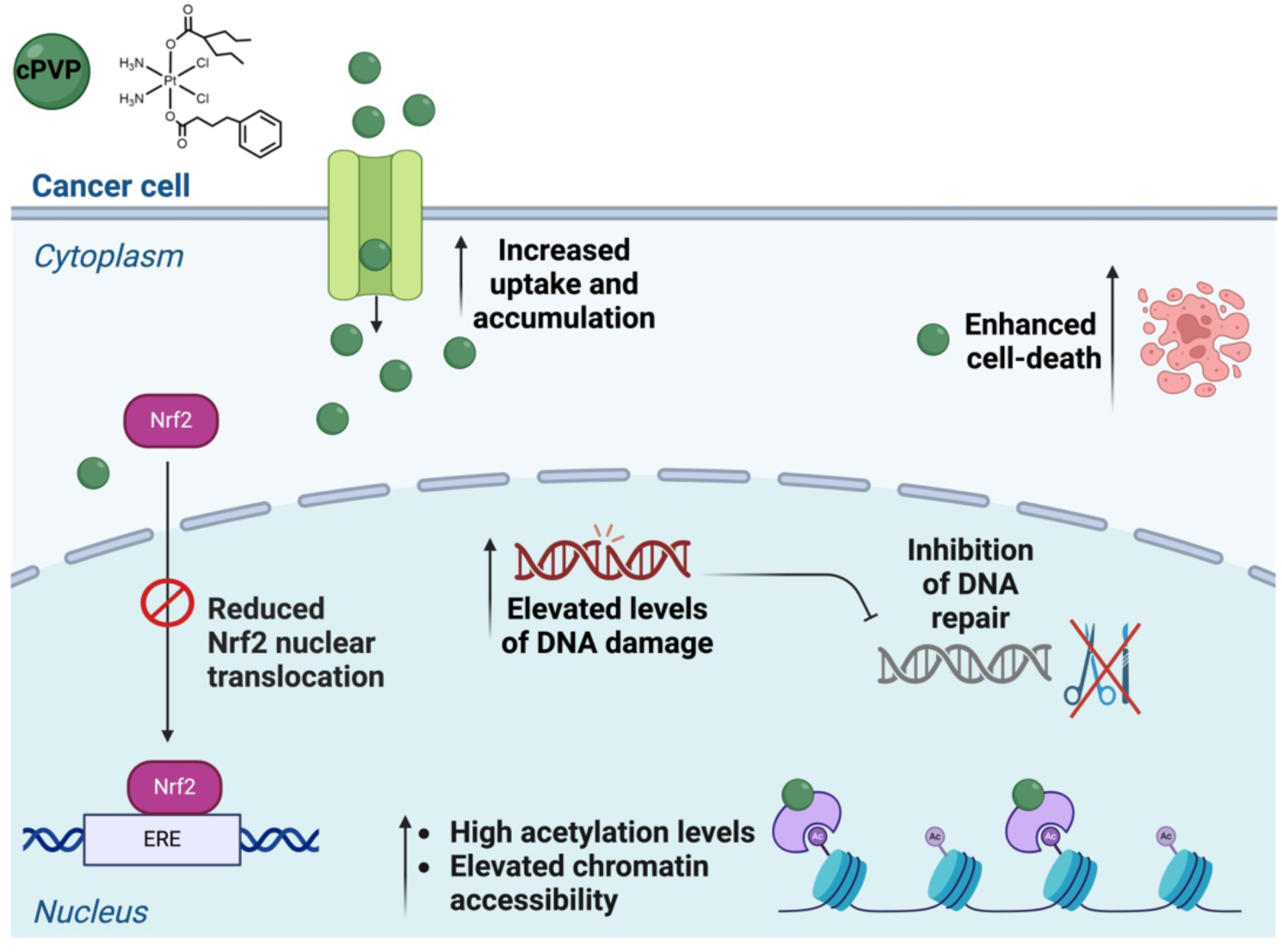
Graphic summary of the multiple simultaneous effects exerted on cancer cells by cPVP.

In conclusion, we’ve synthesized a novel Pt(IV) prodrug that exerts multiple simultaneous detrimental effects on cancer cells, and appears not to induce resistance. Experiments in a mouse mesothelioma model indicate cPVP treatment can increase patients’ survival. The ease of synthesis, low-dose efficiency, and the fact that cPVP’s sub-components (cisplatin, VPA, and PhB) are all FDA approved, position cPVP as a promising chemotherapy and makes it a good candidate for future pre-clinical and clinical studies.

## Materials and methods

### cPVP synthesis

A solution of valproic anhydride (75 mg, 0.28 mmol) dissolved in 1 mL of DMSO (Bio-Lab ltd., Jerusalem, Israel) was added to a suspension of oxoplatin (60 mg, 0.18 mmol) in DMSO (5 mL) and stirred for over-night at room temperature. After confirming the formation of monocarboxylato monohydroxido Pt(IV) by ^195^Pt NMR chemical shift (1037.1 ppm), the clear yellow reaction mixture was extracted several times with a mixture of diethyl ether and petroleum ether mixture (20:1; Bio-Lab ltd., Jerusalem, Israel) to achieve a yellow sticky solid. It was then lyophilized over-night to remove the remaining solvent traces and used in the next step without further purification. The crude solid (71 mg, 0.15 mmol) was mixed with 4-phenylbutyric anhydride (235 mg, 0.76 mmol) following the dissolution in 5 mL of acetonitrile (HPLC grade; J.T. Baker, Phillipsburg, NJ, USA). After stirring for overnight at room temperature, the reaction mixture was injected directly in preparative RP-HPLC (Thermo Scientific UltimaMate 3000 station) equipped with a reverse-phase C18 column (Phenomenex Kinetex, 250 mm x 4.60 mm, 5 μm, 100 Å) and purified with (0–100)% acetonitrile gradient ran over 15 min followed by 15 min at 100% acetonitrile. The pure fractions were combined before lyophilization to afford the desired product as bright yellow flaky solid. Yield 17 mg (20%). RT = 13.5 min (purity >97%). ^195^Pt NMR (107.5 MHz, MeOD) *δ*: 1087.9; ^1^H NMR (500 MHz, MeOD) *δ*: 7.27 – 7.18 (m, 4H, ArH_PhB_), 7.15 (m, 1H, ArH_PhB_), 6.58 (br, NH_3_), 2.66 (t, 2H, *J* = 7.6 Hz, *^γ^*CH_2 PhB_), 2.43 – 2.34 (m, 3H, *^α^*CH_2 PhB_ & CH_Val_), 1.93 – 1.85 (m, 2H, *^β^*CH_2 PhB_), 1.61 – 1.53 (m, 2H, CH_2 Val_), 1.40 – 1.29 (m, 6H, CH_2 Val_), 0.91 (t, 6H, *J* = 7.1 Hz, CH_3 Val_); ^13^C NMR (125 MHz, MeOD) *δ*: 187.4, 184.1, 143.4, 129.7, 129.3, 126.7, 48.6, 36.4, 36.2, 28.9, 21.7, 14.5; ESI-MS calculated for [C_18_H_32_Cl_2_N_2_O_4_Pt + Na]^+^: 628.13, found 629.06; Elemental analysis (%) calculated for C_18_H_32_Cl_2_N_2_O_4_Pt:2.5H_2_O: C 33.18, H 5.72, N 4.30,, found C 32.99, H 5.04, N 4.65. For anhydrides synthesis 50 mM solution of starting free acid in chloroform were stirred overnight in the presence of 0.6 Eq of EDC. The mixture was washed three times with citric acid aqueous solution (1g/100 mL), and three times with sodiumbicarbonate aqueous solution (1g/100 mL). It was then dried on sodium sulphate, and finally evaporated to dryness under reduced pressure.

### Cell culture

A2780 cells (93112519, Sigma-Aldrich, Rehovot, Israel) were cultured in RPMI 1640 Medium (Biological Industries (BI), Beit Haemek, Israel), supplemented with 10% (v/v) fetal bovine serum (FBS; BI), 2 mM L-glutamine (BI), 1 mM sodium pyruvate (BI), 100 U/mL penicillin and 0.1 mg/mL streptomycin (BI). A549 cells (CCL-185^TM^, ATCC, Manassas, VA, USA) were cultured in DMEM (BI) containing 4.5 g/L D-glucose and supplemented with 10% (v/v) FBS, 2 mM L-glutamine, 1 mM sodium pyruvate, 100 U/mL penicillin and 0.1 mg/mL streptomycin. The AB12 cell line was kindly provided by prof. Zvi Friedlander’s lab at the Hadassah Medical Center. AB12 cells were grown in DMEM culture medium (Gibco) supplemented with 10% FBS (BI), 2 mM l-glutamine, penicillin (100 U/ml), streptomycin (100 ug/ml), neomycin (100 ug/ml) and sodium pyruvate (2mm) (BI).

All cultures were maintained in a humidified 5% CO_2_ incubator at 37 °C. Cells were sub-cultured every 3–4 days to maintain logarithmic growth. Cells seeding for all experiments was done 24 hours pre-treatments, allowing cells adherence.

### Cytotoxicity

the cytotoxic effects of cisplatin (Pharmachemie BV, Teva group, Haarlem, The Netherlands) and cPVP against A2780, and A549 tumor cells were evaluated using the well-established MTT [3-(4,5-dimethylthiazol-2-yl)-2,5-diphenyltetrazolium bromide] colorimetric assay (55). In short, cells were seeded into 96-wells plate at a density of 5×10^3^, 2.5×10^3^, and 7×10^3^, respectively, before treatment with ascending concentrations (0-20,000 nM) of cisplatin or cPVP for 72 hours. Media was then replaced with 0.5 mg/mL MTT solution (Sigma-Aldrich, Rehovot, Israel) for one-hour incubation, which was finally replaced with 100 µL/well DMSO. Absorbance was recorded at λ=570 nm and 690 nm, using Cytation 3 Imaging Reader (BioTek, Winooski, VT, USA). Final results were obtained from Δ OD (570–690) nm. IC_50_ values were calculated using the online calculator of AAT Bioquest (56).

### DNA samples preparation for Inductively Coupled Plasma Mass Spectrometry (ICP-MS) analysis

For DNA-damage evaluation, A2780 cells were seeded at 2×10^6^ cells / 60 mm dish and A549 at 2×10^6^ cells / 100 mm dish. Cells were incubated with the indicated doses of cisplatin or cPVP for two hours and then harvested for initial damage levels. For repair assay, the drug containing media was replaced with fresh media for repair-incubation. At the indicated time points, cells were harvested by scraping into ice-cold PBS (BI).

For the assessment of DNA-damage accumulation post-treatment, A2780 cells were seeded at a density of 2×10^6^ cells / 60 mm dish. Cells were treated with 3 µM of cPVP or 20 µM of cisplatin for two hours, followed by treatment removal, wash with PBS, and incubation in fresh media for the indicated times. At each time point cells were harvested by scraping into ice-cold PBS.

In all cases DNA was extracted by using DNeasy® Blood & Tissue kit (69504, Qiagen, Hilden, Germany), and quantified by the QuantiFluor® dsDNA System (E2670, Promega corporation, Madison, WI, USA), following manufacturers’ protocols.

### Cells samples preparation for ICP-MS analysis

To assess cellular accumulation of platinum, A2780 cells were seeded in 6-well plates, at a density of 1×10^6^ cells / well. After the indicated treatments, cells were washed once with PBS and harvested by scrapping into 0.4 mL of fresh PBS.

### ^195^Pt content determination by ICP-MS

Prior to analysis, DNA or cells samples were subjected to acidic digestion. DNA (2 µg) or cells suspension in PBS (0.1 mL) were transferred to scintillation bottles with Silicon/Teflon caps (CSI analytical innovations, Petach-Tikva, Israel) and 1 mL of 70% nitric acid (HNO_3_; Bio-Lab ltd., Jerusalem, Israel) was added. Properly capped samples were placed in a sand-bath and heated for two hours at 90 °C. Then, caps were removed and full evaporation was allowed over-night at the same heat. Samples were re-suspended in 1 mL of 1% HNO_3_, followed by vigorous vortex and shaking. Samples were centrifuged at 10,000 rpm for 10 minutes at room temperature and transferred to ICP-MS suitable Polypropylene scintillation vials (Thermo Fisher Scientific Inc., Waltham, MA USA). Calibration curve ranging from 0 to 10,240 pg/mL was freshly prepared from the commercial cisplatin solution (1 mg/mL) diluted with HPLC-grade water (J.T. Baker, Phillipsburg, NJ, USA), and was subjected to the same procedure described. Samples and calibration curve were measured by Agilent 8900 Triple Quadrupole ICP-MS instrument at the Institute of Earth Sciences core facility of The Hebrew University. ^195^Pt content of DNA samples was normalized to µg of DNA for comparison.

### Damage-seq

A2780 cells were seeded at 5×10^6^ cells per 100mm dish before incubation with 200 µM of cisplatin or 30 µM cPVP for two hours, then harvested by scraping into ice-cold PBS. DNA was extracted and quantified as mentioned above, taking 2 µg of each DNA sample for library preparation. Damage-seq was performed as previously described by Hu et al. (57) without biotin purification after the primer extension step (58). Briefly, Genomic DNA was sheared by sonication with Bioruptor Pico sonicator (Diagenode Inc., Denville, NJ, USA) to generate fragments averaging 300 bp in length. Damaged DNA immunoprecipitation was performed as described previously, with anti-cisplatin modified DNA antibody (ab103261, Abcam, Cambridge, UK). Library quality was assessed using Agilent 4200 TapeStation. Qualified libraries were pooled and sequenced on a NovaSeq 6000 Illumina sequencer. Reads were processed following the steps mentioned in Hu et al. (45, 57) Reads contain Ad1 adapters were discarded by cutadapt (version 3.5) (59)and were aligned to GRCh38 genome using bowtie1 (version 1.3.1) (60). Then, Picard MarkDuplicates (61) (version 2.26.10; http://broadinstitute.github.io/picard accessed on 10 August 2023) was used to remove read duplicates. Next, unique reads in BED format were further filtered with Bedtools (version v2.27.1) (62) and custom BASH scripts. To plot average Damage-seq signal at ATAC-seq peaks, the gene’s annotation file was downloaded from Ensembl, assembly GRCh38, release 96. Only peaks that don’t overlap genes were obtained using custom scripts and Bedtools (version 2.27.1) slop and merge commands. Reads intersecting peak list were obtained using Bedtools intersect command and profiles flanking ATAC-seq peaks were created using the R (version 4.1.3) Bioconductor genomation package (version 1.26.0,) (63). For box plot analysis, read counts for each peak interval were obtained using Bedtools coverage command, and plotted using ggplot2 package (version 3.3.5 (64)). GG coordinates in the entire human genome were extracted with FUZZNUC (version 6.6.0.0) from the EMBOSS package (65). GG profiles were plotted in the same way described for Damage-seq profiles.

### Western-blot

A2780 cells were seeded at density of 1.5 - 3×10^6^ cells / well in 6-well plates, and treated for 24 hours with either 5 µM of cPVP or bisPhB, or 30 µM cisplatin. Cells were lysed under ice-cold conditions, using 50 mM Tris-HCl (pH 7.5), 150 mM NaCl, 1 mM EDTA, 1% Triton X-100 lysis buffer supplemented with protease inhibitors (539134, Calbiochem, San Diego, CA, USA), and collected by scrapping. Lysates were centrifuged at 16,000 rcf, for 10 minutes at 4°C, and protein content was quantified utilizing the Bradford assay as above. Equal amounts of 45 μg protein samples were electrophoresed using 12% TGX stain-free gel (Bio-Rad, Hercules, CA, USA) in TG-SDS running buffer (Bio-Lab ltd., Jerusalem, Israel) and blotted onto PVDF membranes (Trans-Blot Turbo Transfer Pack; Bio-Rad Hercules, CA, USA). The membrane was blocked using 5% (w/v) Difco^TM^ skim milk (232100, BD Life Sciences, Spark, MD, USA) in TBST solution (50mM Tris/Tris-HCl, 150mM NaCl,; Bio-Lab ltd., Jerusalem, Israel. 0.1% Tween® 20 Detergent) for 1h and incubated over-night with primary anti-H3K9Ac antibody at 4°C (C5B11, 9694T, Cell Signaling Technology, Danvers, MA, USA). Membranes were incubated for 1 hour with secondary anti-rabbit HRP-conjugated antibody (NA934, Cytiva, Marlborough, MA, USA) in 5% (w/v) Difco^TM^ skim milk and developed using the ChemiDoc XRS+ imager (Bio-Rad, Hercules, CA, USA).

### ATAC-seq

ATAC-seq was performed as previously published (46) using an ATAC-seq kit (53150, Active motif, Carlsbad, CA, USA) and according to the manufacturer’s protocol on 100,000 cells treated with 30 µM of cPVP or 200 µM of cisplatin for two hours. Prior to beginning the ATAC-seq protocol drug-containing media was replaced with 200 units/mL DNase (LS002006, Worthington Biochemical Corp, Lakewood, NJ, USA) in 1.2 mM MgCl2 and 0.5 mM CaCl2 for 30 minutes. ATAC-seq libraries were purified and size selected with HighPrep PCR beads (AC-60005, MagBio, Gaithersburg, MD, USA). Two rounds of size selection were performed. In the first, 0.6X beads were added, and the supernatant recovered. In the second, 0.6X beads were added to the supernatant and DNA was eluted from the beads in 10 mM Tris-HCl. Library quality was assessed using Agilent 4200 TapeStation. Libraries were pooled and sequenced using an Illumina NextSeq500 sequencer. Sequencing quality was assessed using FastQC (version 0.11.9; https://qubeshub.org/resources/fastqc accessed on 10 August 2023). The adapter sequence was trimmed from each read using Cutadapt (version 1.15) with the command options: -m 10. Reads were aligned to the genome using Bowtie2 (66) with the command options: –very-sensitive -k 10. For each sample we obtained at least 9 million unique aligned reads. Following alignment, reads that were mapped to chromosome Y or mitochondrial chromosome were filtered and PCR duplicates were removed using Picard MarkDuplicates (version 2.26.10 (http://broadinstitute.github.io/picard accessed on 10 August 2023). All bam files of replicates from each condition were merged to create a pooled bam file. Peaks were called separately for each replicate and pooled bam files using macs2 callpeak command (version 2.2.7.1) (67). Peak files from replicates of the same condition were merged using bedtools intersect to create one set of peaks per condition. Peak regions were chosen only if there was more than 50% overlap of the peak region between all replicates per condition. Differential peaks analysis was done using R (version 4.1.3) Bioconductor DiffBind package (version 3.4.11) (68) and plotted using EnhancedVolcano package (version 1.12.0; https://github.com/kevinblighe/EnhancedVolcano).

### Nrf2 nuclear translocation

A2780 cells were seeded at density of 1.5×10^6^ cells / well in 6-well plates. Cells were incubated during 4 hours with 20 and 70 µM of cisplatin, or 3 and 10 µM of cPVP, followed by trypsinization and centrifugation at 600 rcf for 5 minutes at 4°C. Cytosolic and nuclear fractions were then isolated according to the protocol of the Nuclear/Cytosol Fractionation Kit (K266, BioVision, Inc., Mountain View, CA, USA), and protein levels of each fraction were quantified utilizing the Bradford assay (69) with the protein assay dye reagent (Bio-Rad, Hercules, CA, USA) according to manufacturer’s protocol. Proteins from each fraction (5 µg) were loaded on designated 96-well plate of the Nrf2 Transcription Factor Assay Kit (600590, BioVision, Inc., Mountain View, CA, USA) for Nrf2 levels determination, according to the manufacturer protocol. Nrf2 translocation for each treatment was calculated by the ratio of nuclei-to-cytoplasm Nrf2 levels (nuc/cyto ratio) and was normalized to the same ratio of the control sample (UT). Values higher than 1 indicate Nrf2 translocation to the nucleus relative to control.

### Lipid peroxidation

A2780 cells were seeded in 96-well plates at a density of 25×10^3^ cells / well. Cells were incubated for 6 hours with treatments ranging from 0 – 6 µM of cPVP or 0 – 40 µM of cisplatin, before the ROS generating agent 2,2’-Azobis(2-amidinopropane) dihydrochloride (AAPH; final conc. 1 mM in media; 08963, Polysciences, Inc., Warrington, PA, USA) was added for over-night incubation. Lipid peroxidation levels were evaluated by adjusted Thiobarbituric acid reactive substances (TBARS) assay (70). From each well, 50 µL of culture media were transferred to 96-well plate, following by addition of 150 µL 0.57% TBA (Sigma-Aldrich, Rehovot, Israel) in 30% glacial acetic acid (J.T. Baker, Phillipsburg, NJ, USA). Plate was sealed and incubated at 95 °C for 1 hour before being removed to ice for 10 minutes until reaching room temperature. Absorbance at λ=532 nm was immediately measured by the Cytation 3 Imaging Reader. In parallel, MTT assay was conducted on treated cells as described above. Final results were obtained by normalizing TBAR OD to MTT OD for each well.

### XR-seq

A2780 cells were treated for 3 hours with either 30 µM of cPVP or 200 µM of cisplatin. XR-seq was performed as previously described (45, 46). Briefly, 390×10^6^ cells per library were harvested and collected by centrifugation. Cells were resuspended in lysis buffer and lysed using syringes sequentially with 25 and 27 G needles. Low molecular weight DNA was isolated, and samples were incubated with RNase-A (1007885, Qiagen, Hilden, Germany). Primary excision products were pulled down by TFIIH co-immunoprecipitation using anti-p62 and anti-p89 antibodies (sc-293 and sc-292, Santa-Cruz Biotechnology, Dallas, TX, USA). Adapters were ligated on both ends of the excised oligos and immunoprecipitated by the anti-cisplatin antibody. The damages induced by cisplatin were reversed in-vitro by incubating the samples with 200 mM NaCN at 65 °C overnight. DNA was PCR amplified and purified by native polyacrylamide gel electrophoresis (PAGE). Library quality was assessed using Agilent 4200 TapeStation. Libraries were pooled and sequenced on a HiSeq 2500 Illumina sequencer. Quality score for each nucleotide was analyzed using the fastx-toolkit to ensure only high quality reads were processed. The adapter sequence was trimmed from each read using Trimmomatic (71) version 0.36 with the command options: ILLUMINACLIP:adapter_sequence.txt: 2:30:10. Using this adapter sequence: TGGAATTCTCGGGTGCCAAGGAACTCCAGTCACNNNNNNATCTCGTATGCCGTCTTCTG CTTG. Following adapter removal, 50 nt length reads were filtered. Reads were aligned to the genome using Bowtie (version 1.3.1) with the command options: -q –nomaqround –phred33-quals - p 32 -m 4 -n 2 -e 70 -l 20 –chunkmbs 800 –best –S. For each sample we obtained 4.6–7.8 million unique aligned reads. Following alignment, reads mapped to chromosome Y or mitochondrial chromosome were filtered and PCR duplicates were removed using Picard MarkDuplicates (version 2.26.10 (http://broadinstitute.github.io/picard accessed on 10 August 2023). To plot average XR-seq signal along genes, the gene’s annotation file was downloaded from Ensembl, assembly GRCh38, release 96. Non-overlapping regions around the TSS were obtained using custom scripts and Bedtools (version 2.27.1) slop and merge commands. Only non-overlapping genes that were longer than 5000 were included in the plot. Reads intersecting gene list were obtained using Bedtools intersect command and strand-specific profiles over the TSS were created using the R (version 4.1.3) Bioconductor genomation package (version 1.26.0). For box plot analysis, read counts for each gene interval were obtained using Bedtools coverage command, and plotted using ggplot2 package (version 3.3.5). Normalized profiles of XR-seq at ATAC-seq peaks were created as described for Damage-seq profiles. All XR-seq profiles were normalized to Damage-seq profiles.

### Resistance induction

The long-term exposure protocol was based on the work of Behrens et al. (42). Freshly thawed A2780 cells were given one week recovery before determining initial IC_50_ values for cisplatin and cPVP using the MTT assay, as described above. Cells were cultured under three condition: long-term cPVP exposure, long-term cisplatin exposure, and unexposed control. Initial drugs concentrations were 0.003X of the initial IC_50_ (0.02 nM of cPVP and 4 nM of cisplatin), introduced to cells three times for a 72 hours periods during 3 weeks, allowing growth to recover between cycles. Following 3 cycles of exposure, drugs doses were doubled and procedure was repeated until exposure dosage reached 4 times the initial IC_50_ values. With high doses, cycles were extended to 2 weeks each, in order to allow growth recovery before the next exposure. After 36 cycles of exposure, cells were cultured in drug-free media for an additional 4 weeks, and final IC_50_ values were assessed by the MTT assay.

### In-vivo experiments

BALB/c mice were purchased from Envigo. 7 weeks old female mice were used for the experiments. AB12 cells were harvested at 70% confluence and injected (1 × 10^5^) to the peritoneum of BALB/c mice (10 mice per group). cPVP (200 µg) was dissolved in a solution containing 10% cremophor and 10% ethanol in saline and were injected to the peritoneum of tumor bearing mice on days 6 and 13. The control group received the vehicle solution without cPVP. To determine survival, mice were routinely monitored and when they developed significant ascites or their general condition deteriorated, they were euthanized. Tumor tissue was collected from the peritoneal cavity immediately after the mice were anesthetized to confirm tumor development.

### Statistical analysis

All data represents the results from at least two biological replicates performed in at least two technical repetitions. Values are reported as mean ± SEM unless otherwise specified. Statistical significance was calculated by student’s t-test for cisplatin vs. cPVP unless otherwise mentioned. Significance is represented by \**P* < 0.05, \*\**P* < 0.01, and \*\*\**P* < 0.001.

## Ethics approval

In vivo experiments were approved by the Animal Care and Use Committee of the Hebrew University (research # MD-22-16940-5). Patient consent is not applicable.

## Data sharing plan

All raw and processed sequencing data generated in this study have been submitted to the NCBI Gene Expression Omnibus (GEO; https://www.ncbi.nlm.nih.gov/geo/) under accession number GSE243355.

## Competing interests

The authors declare no competing interests.

## Funding

This work was funded by the Israel Science Foundation grants (1710/17 and 482/22) administered by the Israeli Academy for Science and Humanities and the Israel Cancer Association grants (20191630 and 20210078) to SA, the Alex Grass Center for Drug Design and Novel Therapeutics grant administered to RK (3175006046) and the Israel Science Foundation (1002/18) to DG and the Israel Science Foundation (1728/20) to OW. SA is the recipient of the Jacob and Lena Joels memorial fund senior lectureship. RK is affiliated with the Richard and Jean Zarbin Chair in Medical Studies and with the Bloom Center of Pharmacy at The Hebrew University. YNK is the recipient of the Dalia and Eli Hurvitz foundation scholarship for outstanding PhD research students. HGB is the recipient of the Manja Gideon Foundation scholarship in Switzerland and the Lev-Tzion scholarship from the Israel Council for Higher Education. EH is the recipient of the Israel Council for Higher Education Scholarship.

## Author contributions

YNK, SA, RK, and DG conceptualized the project, YNK, ES, HGB, EH, HH, SK, YS, MFZ, OC, and HH performed the investigation, YNK, ES, HGB, EH, HH, SK, YS, MFZ, and HH performed formal analysis, SA, RK, DG and OW supervised the work and acquired funding, YNK and SA wrote the original draft and all authors reviewed and edited the manuscript.

## Acknowledgements

The authors would like to thank Joshua Kahn for technical assistance in the project, Dr. Abed Nasereddin and Dr. Idit Shiff from the Core Research Facility (CRF) at The Faculty of Medicine in The Hebrew University for support in sequencing protocols, Dr. Ofir Tirosh from the Institute of Earth Sciences core facility of The Hebrew University for running ICP-MS samples, and Dr. Avital Parnas for valuable advice and feedback in manuscript preparation. Figure artwork was generated using biorender.com and graphs were generated using GraphPad prism version 8.0.1.

## Notes

**Competing Interest Statement:** The authors declare no potential conflict of interest.

### Competing Interest Statement

The authors have declared no competing interest.

