## Supplementary Figures for "A Novel Cisplatin-Based Prodrug Inhibits Lysine Deacetylases, Suppresses Nucleotide Excision Repair, and Overcomes Resistance"

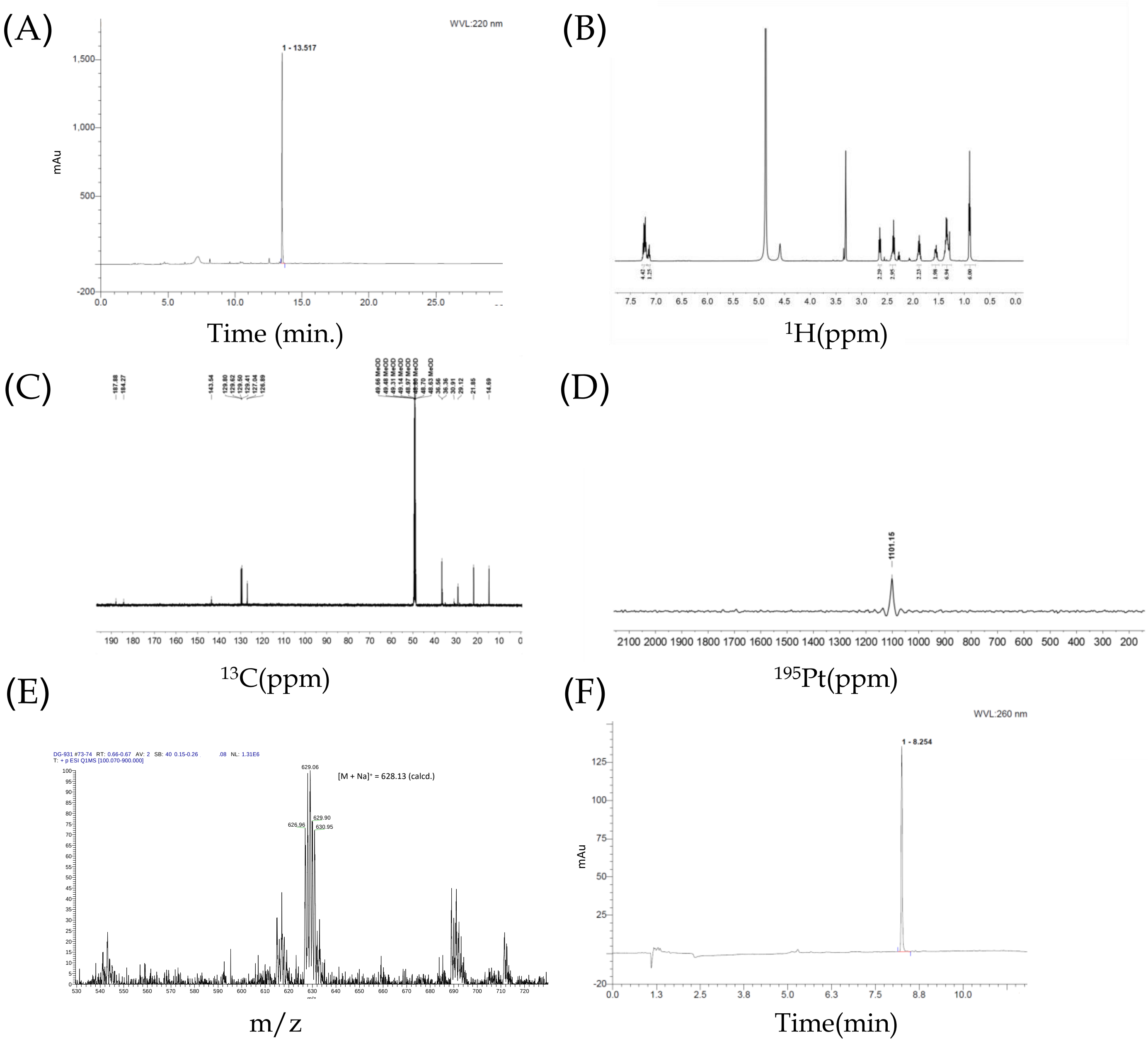


**Figure S1. Full characterization of cPVP. (A)** HPLC chromatogram of cPVP ran with (0 – 100)% acetonitrile gradient over 15 min followed by 15 min at 100% acetonitrile, in a reverse-phase C18 column (250 mm x 4.60 mm, 5 µm, 100 Å). **(B)** 1H NMR of cPVP in MeOD. **(C)** 13C NMR of cPVP in MeOD. **(D)** 195Pt NMR of cPVP in MeOD. **(E)** ESI-MS (+ve) data of cPVP. **(F)** HPLC chromatogram of cPVP after three years of storage at -20□C. Ran with (0 – 100)% acetonitrile gradient over 25 min followed by 5 min at 100% acetonitrile, in a reverse-phase C18 column (100 mm x 4.60 mm, 2.6 µm, 100 Å).

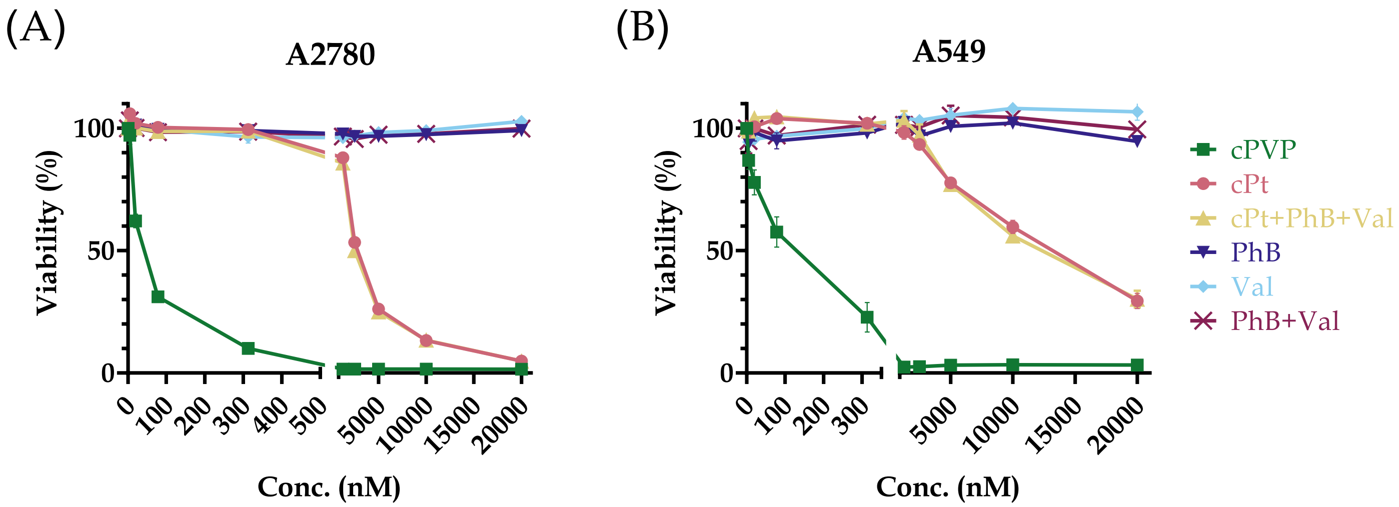


**Figure S2. Co-administration of cisplatin with inhibitors. (A)** Cytotoxicity of cPt, cPVP and its' components when co-administered in A2780 cells. Cells were incubated for 72 hours with a range of concentrations of either cPVP, cPt, cPt+PhB+VPA (in a 1:1:1 ratio), PhB, VPA, or PhB+VPA (in a 1:1 ratio), followed by MTT assay. Data presented as percent of viability normalized to untreated control. **(B)** Same as A except data presented for A549 cells. Graphs present mean ± SEM. Results are based on at least two biological replicates performed in quadruplets. ****P<0.0001 for all treatments vs. cPVP based on two-way ANOVA for both cell-lines.

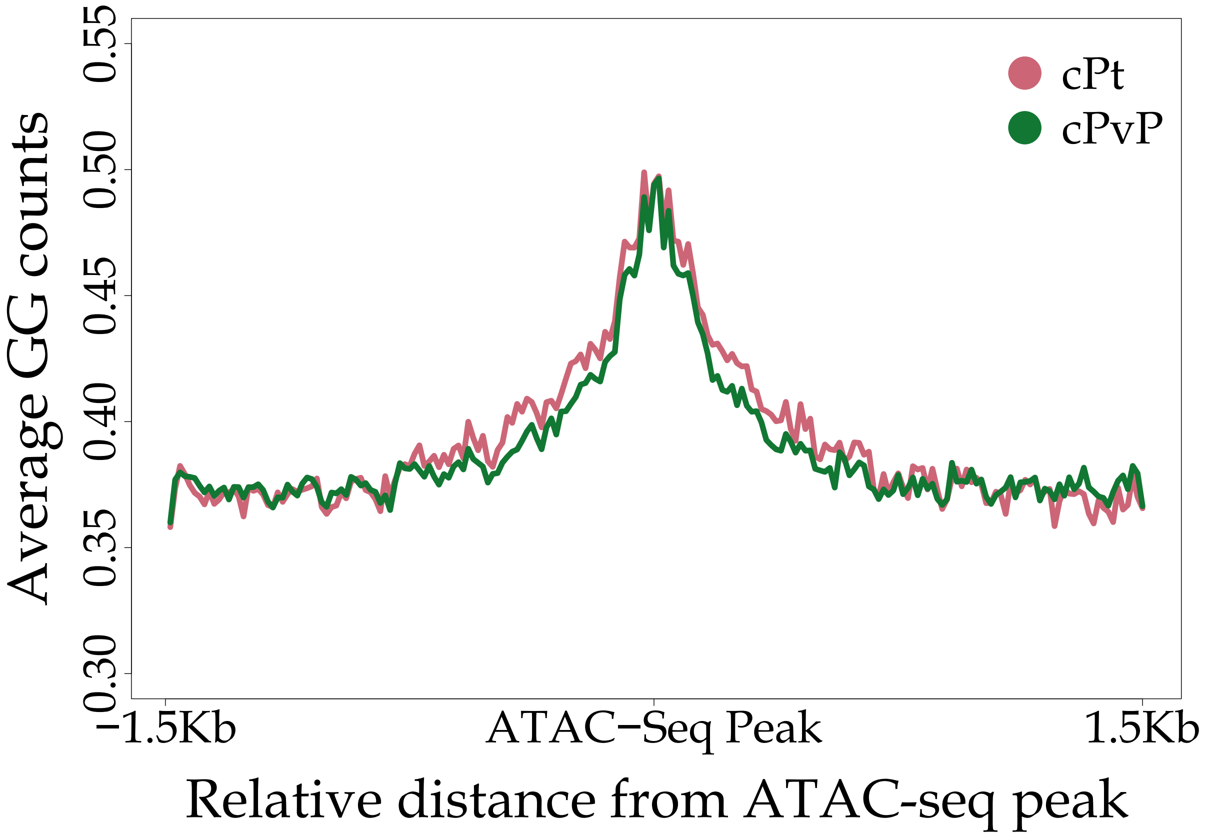


**Figure S3. GG profile of A2780 cells.** Average GG frequency at ATAC-Seq peaks regions measured following cPt and cPVP treatments in A2780 cells. GG frequency is plotted at ATAC-Seq peaks and 1.5 Kb flanking regions with a bin size of 15 nt. Shadow represent 95% confidence interval for the mean.

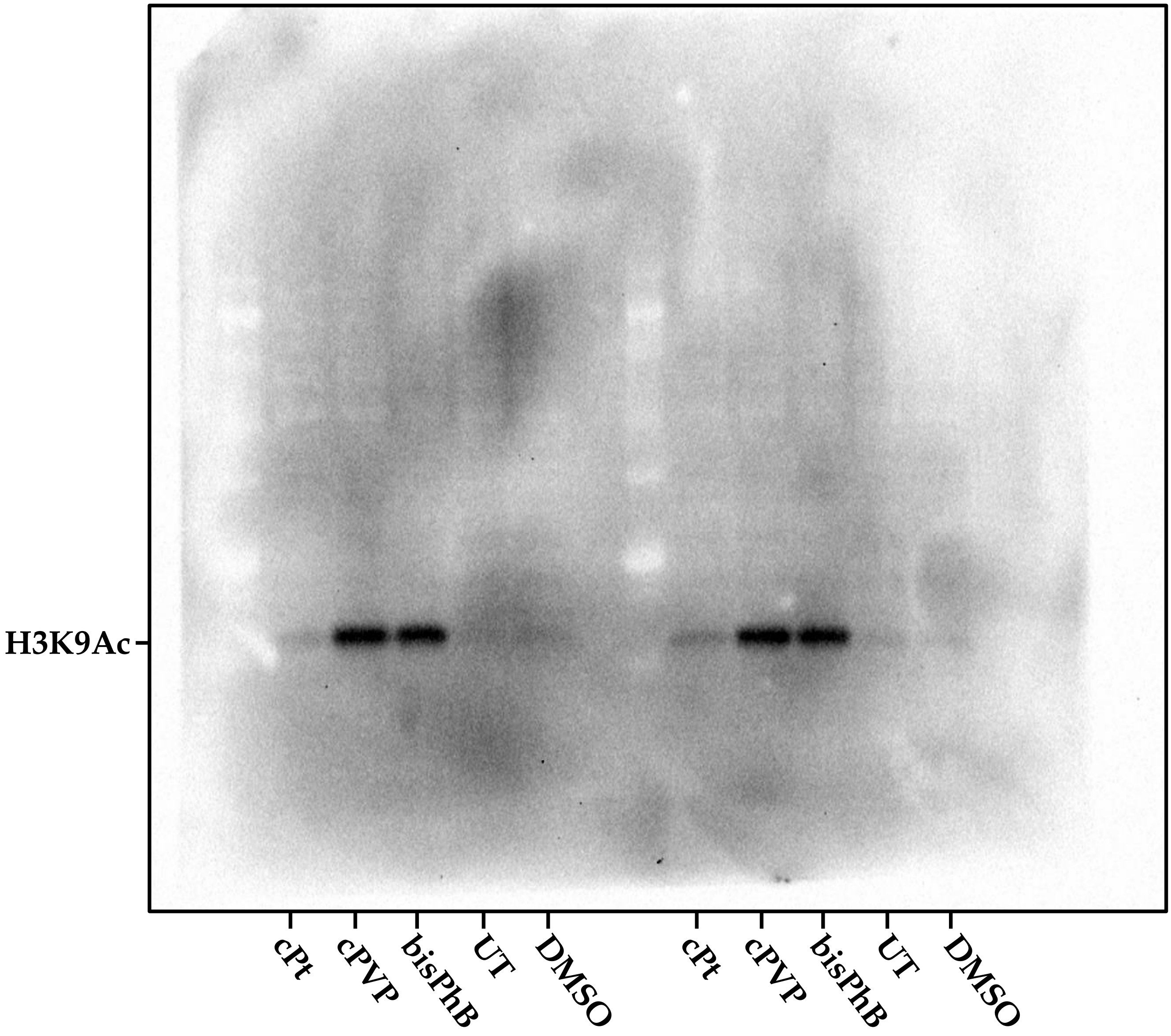


**Figure S4. Full western-blot image corresponding to Figure 4A.** Histone H3 acetylation analysis using H3K9Ac antibody. A2780 cells were treated with 5 μM of cPVP, 5 μM of bisPhB or 30 μM of cPt for 24 hours.

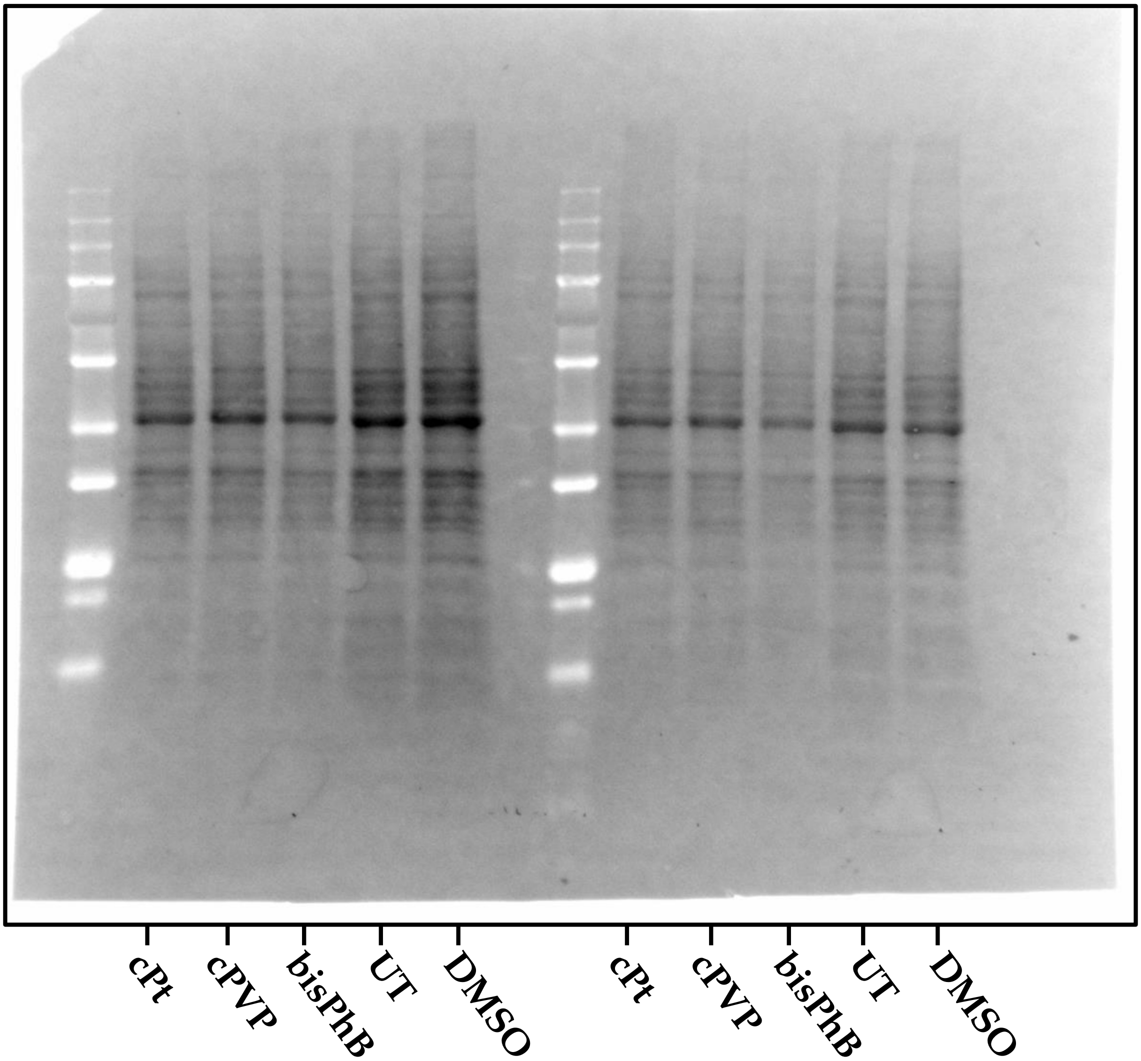


**Figure S5. Total protein membrane image used to quantify Western blot in Figure 4A.**

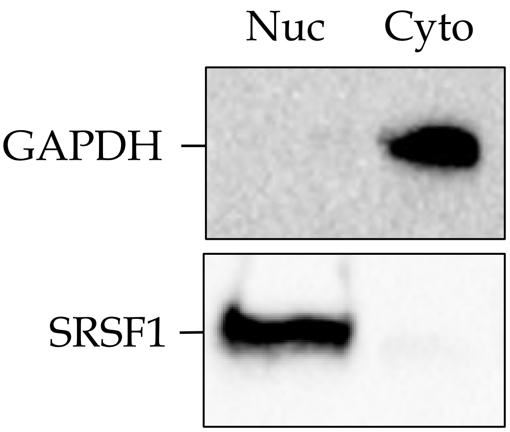


**Figure S6. Cytosolic and nuclear fractions analysis.** Western-blot for the cytoplasmic marker GAPDH and nucleic marker SRSF1 in nuclear and cytosolic fractions from A2780 cells.
